# ATP-sensitive K^+^ channels control the spontaneous firing of a glycinergic interneuron in the auditory brainstem

**DOI:** 10.1101/2020.06.17.155671

**Authors:** Paulo S. Strazza, Daniela V.F. de Siqueira, Nikollas M. Benites, Ricardo M. Leão

## Abstract

Cartwheel neurons from the dorsal cochlear nucleus (DCN) are glycinergic interneurons and the primary source of inhibition on the fusiform neurons, the principal excitatory neuron in the DCN. Most cartwheel neurons present spontaneous firing (*active* neurons), producing a steady inhibitory tone on fusiform neurons. In contrast, a smaller fraction does not fire spontaneously (*quiet* neurons). Additionally, hyperactivity of fusiform neurons is seen in animals with behavioral evidence of tinnitus. Due to its relevance in controlling the excitability of fusiform neurons, we investigated the ion channels responsible for the spontaneous firing of cartwheel neurons. We found that quiet neurons express an outward conductance not seen in active neurons, which generates a stable resting potential. This current was sensitive to tolbutamide, an ATP-sensitive potassium channel (K_ATP_) antagonist. After its inhibition, quiet neurons start to fire spontaneously, while the behavior of active neurons was not affected. On the other hand, in active neurons, K_ATP_ agonist diazoxide activated a conductance similar to the K_ATP_ conductance of quiet neurons and stopped spontaneous firing. According to the effect of K_ATP_ channels on CW neuron firing, glycinergic neurotransmission in DCN was increased by tolbutamide and decreased by diazoxide. Finally, slices incubated with the tinnitus-inducing agent sodium salicylate presented more quiet neurons expressing the K_ATP_ conductance, which increased the proportion of quiet neurons. Our results reveal an unexpected role of K_ATP_ channels in controlling the spontaneous firing of neurons. Additionally, changes in K_ATP_ channel activity of cartwheel neurons can be related to the DCN hyperactivity seen in tinnitus.

## Introduction

Most neurons present a stable resting membrane potential, firing only in response to suprathreshold synaptic inputs. However, several neurons exhibit unstable membrane potentials leading to spontaneous action potential (AP) firing. These neurons express diverse combinations of conductances that drive the membrane potential to values above the AP threshold to initiate spontaneous action potential generation. For instance, thalamic neurons rely on the cooperation of low voltage-activated T-type calcium channels (ICa_T_) and hyperpolarization-activated cationic currents (I_h_) to depolarize the membrane (McCornick and Huguenard, 1992) while the activity of midbrain dopaminergic neurons is not dependent of I_h_ (Mercuri et al., 1995) but mainly on voltage-dependent calcium channels (Grace and Onn, 1989; Puopolo et al., 2007; Putzier et al., 2009). On the other hand, many neuronal types use tetrodotoxin (TTX)-sensitive subthreshold or background (leak) Na^+^ permeable conductances to depolarize the membrane and start spontaneous firing (Leão et al., 2012; Ceballos et al., 2016; Raman et al., 2000; Pennartz et al., 1997; Khaliq and Bean, 2010).

The dorsal cochlear nucleus (DCN) is a subdivision of the cochlear nucleus with a cerebellum-like organization that integrates acoustic information, via the auditory nerve terminals, with multimodal information from other sensory areas and superior auditory stations, conveyed by the parallel fiber system (Oertel and Young, 2004). Hyperactivity of DCN neurons is implicated in the generation of tinnitus (the chronic perception of ringing in the ears) in experimental animal models (Kaltenbach and Afman., 2000; Brozoski et al., 2002) and ablation of the DCN prevents tinnitus induction in animals (Brozoski et al., 2012). The principal neuron in the DCN is the fusiform (also called pyramidal) neuron. It is the main output to the inferior colliculus, integrating the information coming from the auditory nerve with the somatosensory pathways of granule cell axons forming the parallel fibers. Several fusiform neurons present spontaneous firing (*active*), both *in vivo* and *in vitro* recordings, while another large fraction is silent (*quiet*) (Hirsch and Oertel, 1988; Manis, 1990; Parham et al., 1992 Hancock and Voigt, 2002; Leão et al., 2012). The primary determinant of this complex behavior of DCN fusiform neurons is the differential expression of an inwardly rectifying potassium current (I_Kir_). Quiet fusiform neurons express a more robust I_Kir_, which sets the RMP below an activity threshold created by the persistent sodium current (I_NaP_). In contrast, in active neurons, the small magnitude of I_Kir_ sets the RMP above the activity threshold (Leão et al., 2012).

The fusiform neuron receives substantial inhibitory input from glycinergic cartwheel (CW) interneurons, which are also innervated by the parallel fibers, forming a feed-forward inhibitory circuit on the fusiform neuron that controls its excitability during the parallel fiber activity (Golding and Oertel, 1997). CW neurons present a complex and dynamic firing behavior, distinguished by different patterns of bursting and tonic firing, and the presence of clusters of high-frequency action potentials riding on the top of a slow depolarization, created by the activation of voltage-dependent sodium and calcium channels, called complex spikes (Golding and Oertel, 1997; Kim and Trussell, 2007, Bender et al., 2012). Like fusiform neurons, CW neurons also present *quiet* and *active* behavior. While most CW neurons fire action potentials spontaneously, both simple and complex, in a variety of firing patterns (Kim and Trussell, 2007; Zugaib et al., 2016). However, a small fraction of CW neurons present a stable resting membrane potential and do not, or seldom, fire spontaneous action potentials (Kim and Trussell, 2007; Zugaib et al., 2016). Nevertheless, it remains unknown which ionic mechanisms determine whether a CW neuron presents active or quiet behavior. Interestingly, incubation slices with high concentrations of sodium salicylate, a tinnitus-inducing agent, increases the proportion of quiet CW neurons (Zugaib et al., 2016), and this effect reduces inhibition of fusiform neurons, which could be relevant for the generation of tinnitus by high concentrations of salicylate.

Because the decrease in the spontaneous firing of CW neurons can have a marked effect on the inhibition of fusiform neurons and could be relevant for tinnitus generation, we investigated the ionic conductances governing spontaneous firing of DCN CW neurons. We found an unexpected role for ATP sensitive potassium (K_ATP_) channels in the control of firing of CW neurons. Additionally, we found that the tinnitus-inducing agent salicylate increases the number of CW neurons expressing K_ATP_ currents, which increases the proportion of quiet CW neurons, decreasing the inhibition on the Fusiform neurons, which can be a mechanism generating hyperactivity in these neurons and contribute for tinnitus generation.

## Methods

### Animals and preparation of slices

Male Wistar rats (p18-22) were obtained from the Central Animal Facility of the School of Medicine of Ribeirão Preto. All protocols were approved by the Ethics Committee on Animal Experimentation (CEUA) under the protocol 007/2017. The animals were killed by decapitation after isoflurane anesthesia, and the brains removed and submerged in an ice-cold solution containing (in mM): NaCl (87), NaHCO_3_ (25), KCl (2.5), NaH_2_PO_4_, (1.25), CaCl_2_ (0.2), MgCl_2_ (7), glucose (25), sucrose (75), 335 mOsm/kgH_2_O, pH 7.4 when bubbled with carbogen (95% O_2_ and 5% CO_2_). Before removing the brain from the skull, the vestibulocochlear nerve (VIII cranial nerve) was cut to prevent damage to the DCN. Coronal slices (200 μm thick) containing the DCN were obtained on a vibratome (Vibratome 1000 Plus) and incubated at 35°C for 45 minutes and subsequently at room temperature in recording solution (artificial cerebrospinal fluid - aCSF), whose composition (in mM) was: NaCl (125), KCl (2.5), NaHCO_3_ (25), NaH_2_PO_4_ (1.25), glucose (10), CaCl_2_ (2), MgCl_2_ (1), 305 mOsm/kgH_2_O, pH 7.4 when bubbled with carbogen (95% O_2_ and 5% CO_2_). In some experiments, 5 mM glucose was used, with 2.5 mM NaCl added to maintain osmolarity. 1 µM of strychnine (STR), 20 μM of picrotoxin was added to the aCSF to inhibit spontaneous glycinergic and GABAergic transmission, respectively. 6,7-dinitroquinoxaline-2,3-dione (DNQX) was added in some experiments to avoid action potential firing from the less frequent spontaneous glutamatergic synaptic currents.

For testing the effect of salicylate, sodium salicylate (1.4 mM) was added to the aCSF and incubated with the slices after cutting and in the perfusion solution, being present throughout the whole experiment, as described previously (Zugaib et al., 2016).

### Electrophysiology

For electrophysiological recordings, the slices were transferred to a perfusion chamber and fixed with a nylon thread mounted in a platinum frame. Neurons were visualized on a microscope (Olympus BX51WI) equipped with IR-DIC optics and an infrared-sensitive camera (Hamamatsu). The slices were continuously perfused (1 ml/min), with aCSF at 33-35°C with an inline heater (Warner Instruments).

Electrophysiological recordings were performed in whole-cell patch-clamp using an EPC-10 amplifier (HEKA Elektronik) or with a Multiclamp700B connected to a Digidata 1440A AD/DA converter board (Molecular Devices), in voltage-clamp and current-clamp mode. For this, borosilicate recording electrodes (BF-150-86-10, Sutter Instruments) were pulled with a horizontal puller (P87-Sutter instruments) and backfilled with an internal solution containing (in mM): potassium gluconate (130), EGTA (0.1), HEPES (10), KCl (20), ATP-Mg (2), GTP-Na (0.2), phosphocreatine-Na (10); 305-310 mOsm/kgH_2_O, pH 7.3 adjusted with KOH, resulting in a tip resistance of 3 to 6 MΩ. Cartwheel neurons were identified by their location in the molecular layer and by the presence of compound action potentials (complex spikes) (Kim and Trussel, 2007). Fusiform neurons were identified by their location in the fusiform layer of the DCN, large cell body, and their regular firing pattern when depolarized with no or very little accommodation (Zhang and Oertel, 1994). Only cells that presented at least one complex spike during a train of action potentials were classified as CW cells and included in analysis. Most recordings were from cells chosen in an unbiased way, but because quiet CW neurons were infrequent (Kim and Trussel, 2007; Zugaib et al., 2016; see Results), many times, we chose quiet neurons by performing cell-attached recordings before entering whole-cell. In this protocol, we considered an active CW neuron when we detected complex spikes or burst firing (Kim and Trussell, 2007).

Voltage-current (VI) relationships were performed by injecting 1000 ms current steps ranging from −300 to 400 pA with 50 pA increments from a membrane potential adjusted to ≅ −80 mV (by applying current as necessary), with intervals between stimuli of 1.5 second. In voltage-clamp experiments, neurons were held at a potential of −65 mV, except when mentioned otherwise. For IV relationships, 4 seconds voltage steps from −115 mV to −25 mV, in 10 mV increments, were applied with an interval of 1.5 seconds between them. Ramp protocols were applied from −100 mV to −50 mV. Ramp duration was 4 seconds (12.5 mV/second).

Spontaneous glycinergic neurotransmission was recorded in the fusiform neurons held at −70 mV in the presence of the glutamatergic AMPA/kainite receptor 6,7-dinitroquinoxaline-2,3-dione (DNQX; 10 μM) and the GABA_A_ antagonist picrotoxin (100 μM). After the experiment, the glycinergic nature of the sIPSCs was confirmed by applying the antagonist of glycine receptors strychnine (10 μM), which completely inhibited the sIPSCs.

The recordings were initiated 1 minute after accessing the whole-cell configuration. Selective ion channel antagonists/agonists were perfused for 3 to 5 minutes before recordings were started in the treatment condition, and then for more 10 to 15 minutes. Uncompensated series resistance was < 20 MΩ, and compensation was in the range of 50 to 70%. Data were low-passed (Bessel 8-pole) at 3 kHz and digitized at 10 kHz.

### Chemicals

All salts were of reagent grade. All solutions were prepared with ultrapure water. DNQX, picrotoxin, strychnine, riluzole, sodium salicylate, tolbutamide, and diazoxide were from Sigma (St. Louis, MO), ZD7288 from Tocris (Bristol, UK), tetrodotoxin from Alomone Labs (Jerusalem, Israel).

### Data analysis

Data were analyzed using custom-made routines in IGOR Pro 6 (WaveMetrics). All membrane potential data was adjusted for a liquid junction potential of −10 mV. As most CW neurons have spontaneous firing, we adopt the modal membrane potential (MMP) as the measure of the intrinsic or resting membrane potential. The MMP was obtained from recordings of the membrane potential of 52 seconds with no current injected. An all-points histogram of the membrane potential with bins of 1 mV was performed, and resting membrane potential was defined as the peak of the histogram (the statistical mode). Neurons with a tonic pattern of firing or quiet neurons generated a histogram with a single peak (Figure 1A and B*i*). In contrast, neurons with more complex firing behaviors generated broader histograms (irregular firing pattern) or histograms with 2 peaks (e.g., regular bursting firing) (Figure 1B*ii*). In this last case, we chose the taller peak as the representative of the MMP.

**Figure 1:**
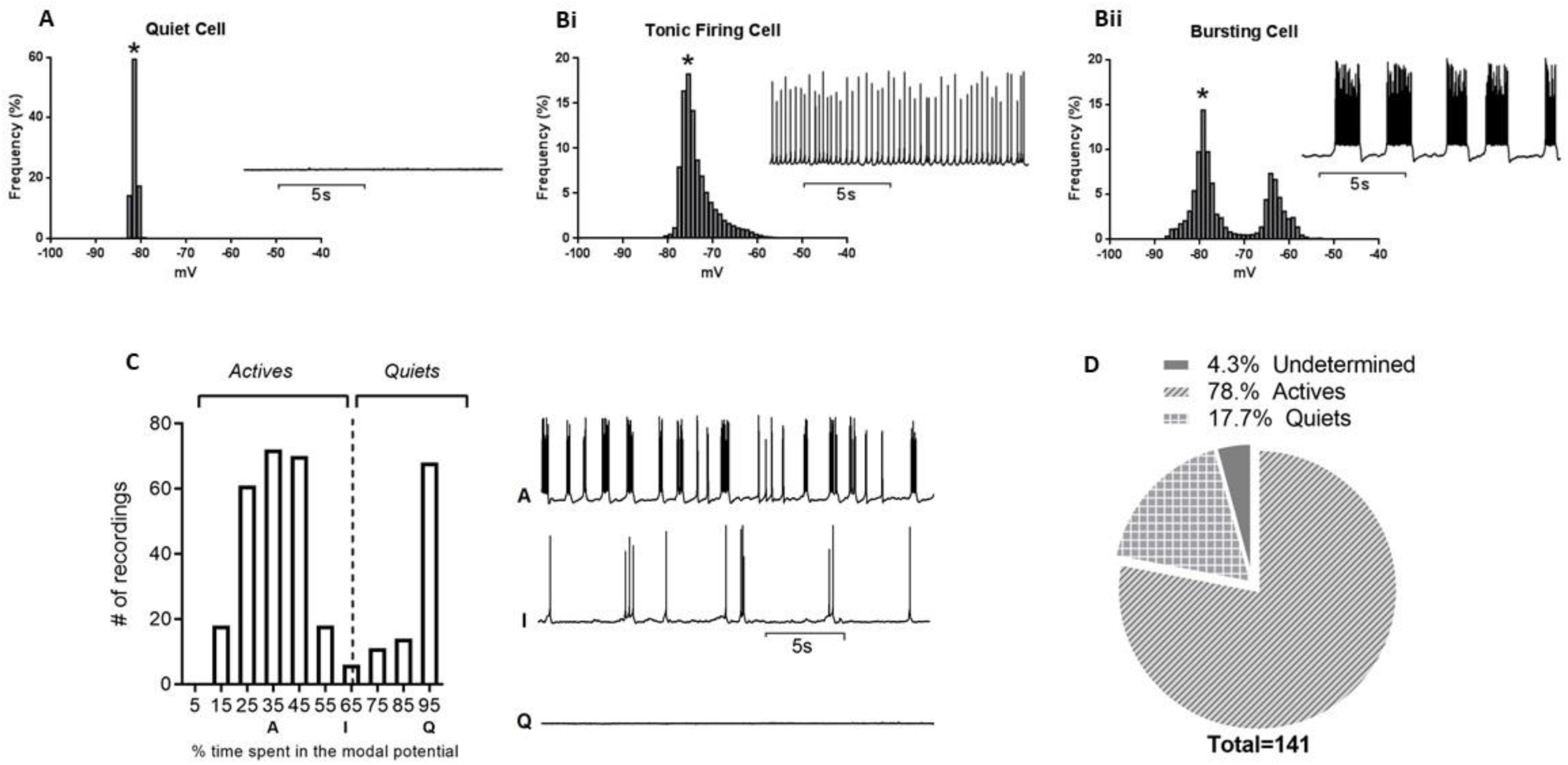
Quiet and active cartwheel neurons. A. histogram of membrane potentials of a quiet neuron (inset). B. histogram of membrane potentials of a tonic active neuron (i, inset) and a bursting active neuron (ii, inset). The stars show the modal membrane potential (MMP). C, histogram showing the percentage of time spent in the modal potential (MMP_%_) of 338 membrane potential recordings of CW neurons. Q= quiet (70%< MMP_%_); I= undetermined (60%< MMP_%_<70%); A=active (MMP_%_<60%). The insets show representative recordings from each type. D. pie chart showing the proportions of active, quiet, and undetermined neurons (data from whole-cell recordings).

Membrane resistance was measured in the current-clamp mode as the slope of linear regression (4 points) of the membrane potential of the three first hyperpolarization pulses of the voltage-current (VI) plot. For this measurement, we adjusted the membrane potential to around −80 mV. The membrane time constant was measured by fitting a single exponential function to the decaying phase of the membrane potential in response to the first hyperpolarization of the VI plot. The depolarization sag was measured as the difference between the peak hyperpolarization and the membrane potential at the last 250 ms of the second hyperpolarizing pulse. The activity threshold was determined as in Leão et al. (2012) as the membrane potential above which neurons start to fire spontaneous action potentials in response to depolarization pulses, starting from a membrane potential around −80 mV.

The slope membrane conductance was measured in voltage-clamp as the slope of the linear regression of 3 points in the IV relationship. The chord conductance was obtained from a regression from the point of maximum current (or from a specific voltage, mentioned) in the IV relationship to the measured or estimated reversal potential of the current.

sIPSCs were analyzed using the “template” model in the software Clampfit (Molecular Devices). 2 to 3 stretches of 1 second were analyzed. For control events, we chose stretches immediately before the drug application, and 5-10 minutes after drug application. sIPSCs were analyzed by their inter-event intervals and amplitudes. Event frequency was calculated using the inter-event intervals. The events detected were checked individually.

Data are shown as the mean ± SEM, except proportions, which are reported as absolute values. Means were compared using two-tailed paired and unpaired Student’s t-tests. Proportions were compared with two-sided Fisher’s exact test. Prism 8.0 (GraphPad) was used for statistical analysis. In the figures *=p<0.05;**=p<0.01; ***=p<0.005; ****=p<0.0001.

## Results

### Active and quiet cartwheel neurons: classification, proportions, passive membrane properties and subthreshold currents

Most CW neurons fire spontaneously at rest, and a small fraction maintains a stable resting membrane potential. However, some CW neurons switch between silent and active states or have very low levels of spontaneous activity (firing rate < 0.5Hz) or episodic short bursts, which makes their classification in two discrete categories more challenging. We, therefore, based our classification of quiet and active neurons on the percentage of time spent around the modal membrane potential and the IV relationships of each neuron. From histograms of the membrane potential variation (Figure 1A-B), we calculated the percentage of time spent in a window of ± 1.5 mV around the modal membrane potential (MMP) of each neuron and compared it with its IV relationship. Quiet neurons had a clear zero current crossing in their IV relationships. They presented more stable membrane potentials, spending from 70% to 100% of the time at MMP, firing none (the majority of quiet neurons) or episodic action potentials during the recording period (Figure 1C). In contrast, active neurons did not present a zero crossing in their IV relationship and spent less than 60% of the time at MMP (Figure 1C; see Figure 2D for the difference between the IV relationships of quiet and active neurons). We observed that 21% of recorded neurons switched from quiet to active, and 9% switched from active to quiet. When such transitions happened, we recorded the membrane potential for four additional minutes and chose the dominant phenotype for analysis. We also observed some few neurons (n=6) that presented frequent state transitions and changes in the levels of spontaneous activity. These neurons spent between 60 to 70% of the time in the MMP (Figure 1C) and presented inconsistent IV relationships, which in some cases, crossed zero and, in other cases, did not. We classified these cells as undetermined, but because of this unsteady behavior, we excluded them from our analysis.

**Figure 2:**
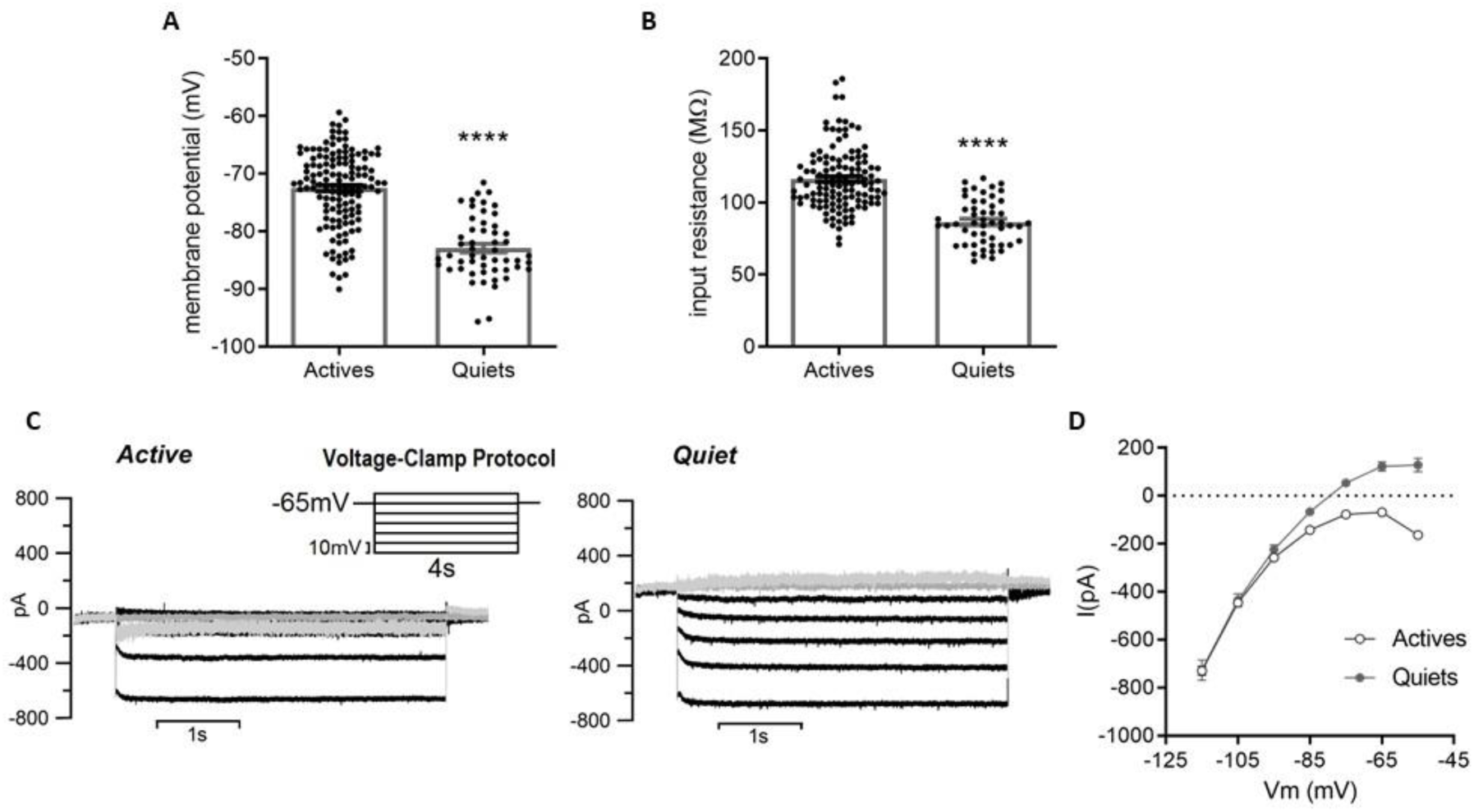
Active and quiet cartwheel neurons presented distinct membrane passive properties and subthreshold currents. A. membrane potential of quiet and active neurons. B. membrane input resistance of quiet and active neurons. C. subthreshold currents of a representative quiet and active cartwheel neuron (in the inset the voltage protocol used; grey trace = −65mV; light grey = −55mV). D. IV relationships of subthreshold currents of cartwheel quiet and active neurons. ****p<0.0001.

From an unbiased sample of neurons (neurons we did nor pre-select before whole-cell; see methods), we found that in the DCN of rats, 78% of CW neurons were active, 17.7% were considered quiet, and 4.3% were considered undetermined (Figure 1D; n= 141). Excluding the undetermined cells, this fraction changes to 81.5% of active cells and 18.5% of quiet cells (n=135). When we included data from cell-attached recordings, this proportion remained similar (82.3% active neurons and 17.7% quiet neurons; n=294). Thus, the prevalence of the active state in CW neurons does not seem to be caused by artifacts from the whole-cell configuration. This proportion was similar to previous reports in rat DCN (Zugaib et al., 2016) and relatively close to the values reported in perforated patch recordings in mouse DCN (Kim and Trussell, 2007).

Next, we compared the passive membrane properties of quiet and active CW neurons. The membrane potential of quiet neurons was −82.9 ± 0.7 mV (n=52), whereas active neurons had a significantly more depolarized membrane potential of −72.5 ± 0.6mV (n=133; p<0,001; Figure 2A). Membrane input resistance was significantly higher in active neurons (Quiet: 86.5 ± 2.2 MΩ, n=52; Active: 116.3 ± 1.9 MΩ, n=123; p<0.001, Figure 2A). Membrane time constant was, accordingly, higher in active neurons (Quiet: 13.1 ± 0.5 ms, n=52; Active: 21.6 ± 0.5ms, n=121; p<0.001). Additionally, active CW neurons presented a more prominent depolarization sag when hyperpolarized (Quiet: 1.3 ± 0.1mV, n=52; Active: 2.1 ± 0.1mV, n=120; p<0.001). These results indicated differences in the expression of subthreshold membrane conductances between quiet and active DCN CW neurons that could account for the generation of the two phenotypes.

Next, we measured the membrane conductances in the sub and near-threshold potentials of quiet and active CW neurons. We found that the IV relationships of quiet and active neurons were similar in the range between −115 to −95 mV (Figure 2C-D) but diverged for more depolarized membrane potentials. Quiet CW neurons presented an IV relationship that crossed the zero current line, a characteristic of neurons with a stable resting membrane potential. In contrast, the IV relationship for active neurons started to move away from zero current around −75 mV, and then became increasingly negative (Figure 2C,D). While the presence of an inward current in near-threshold potentials in quiet neurons is not clear in the IV relationship shown in figure 2D, we observed the development of an inward current around −50 mV in IVs in which we recorded more positive potentials (not shown). These results demonstrated that the membrane conductances of quiet and active CW neurons are distinct, reaching a point of electrochemical equilibrium (the zero current point) in quiet neurons, and deviating from this point in active neurons.

### Persistent sodium current (I_NaP_) drives action potential firing in active and quiet cartwheel neurons

In active DCN fusiform neurons, persistent sodium current (I_NaP_) produces a deviation from the membrane electrochemical equilibrium, generating an inward current that triggers spontaneous firing (Leao et al., 2012). I_NaP_ consistently decreases the threshold for spontaneous firing, called activity threshold (Leao et al., 2012), and blocking I_NaP_ with riluzole increases the activity threshold and abolished spontaneous firing in active fusiform neurons (Leao et al., 2012). Kim and Trussel (2007) described a similar persistent sodium current in mouse CW neurons but did not investigate its role in driving spontaneous firing.

We measured activity threshold in quiet and active CW neurons and found that this parameter is more hyperpolarized in active CW neurons than in quiet neurons (Quiet: −75.9 ± 0.9mV, n=52; Active: −79.9 ± 0.4mV, n=124; p<0.0001; Figure 3A). Accordingly, the activity threshold of quiet neurons was above (more positive) their modal membrane potential (by 7.0 ± 0.9 mV, n=52. Figure 3B), while in active neurons the activity threshold was more negative than their modal membrane potential (by −7.3 ± 0.5 mV, n=124; Figure 3B). To test if I_NaP_ was responsible for the spontaneous firing, we applied riluzole (10 μM), an inhibitor of the persistent component of the sodium current (Leão et al., 2012). Riluzole prevents firing of spontaneous fast Na^+^ spikes in most active neurons (9 out of 12). Of these neurons, riluzole was able to produce stable membrane potentials with no action potential firing in two neurons (Figure 3A), while in the 7 other neurons membrane potential in the presence of riluzole presented oscillations, probably calcium spikes (Kim and Trussell, 2007), which occasionally drove sodium action potentials. All these neurons were still able to fire action potentials when depolarized by injected current (Figure 3E), showing that riluzole did not affect substantially fast-inactivating sodium channels. Interestingly, three neurons continued to fire action potentials tonically after riluzole, suggesting other inward currents may contribute to spontaneous action potentials. Additionally, after the inhibition of spontaneous action potential firing by riluzole or TTX (0,5 μM) the resting membrane potential of active neurons was more depolarized than the membrane potential measured during action potential firing (from −73,0 ± 1,2mV to −60,9 ± 2,7mV; n=17; p=0,0003; Figure 3C,D) suggesting that the after-hyperpolarization (AHP) strongly affects the MMP. On the other hand, in quiet neurons, we observed a small hyperpolarization after riluzole or TTX (from −82.5 ± 2.1 mV to −85.3 ± 2.1 mV, n=7, p=0. 003; Figure 3D), showing that I_NaP_ is activated at resting in CW neurons. After inhibition of spontaneous firing, the resting membrane potential of active neurons was more depolarized than the resting membrane potential of quiet neurons (p<0.0001; Figure 3D).

**Figure 3:**
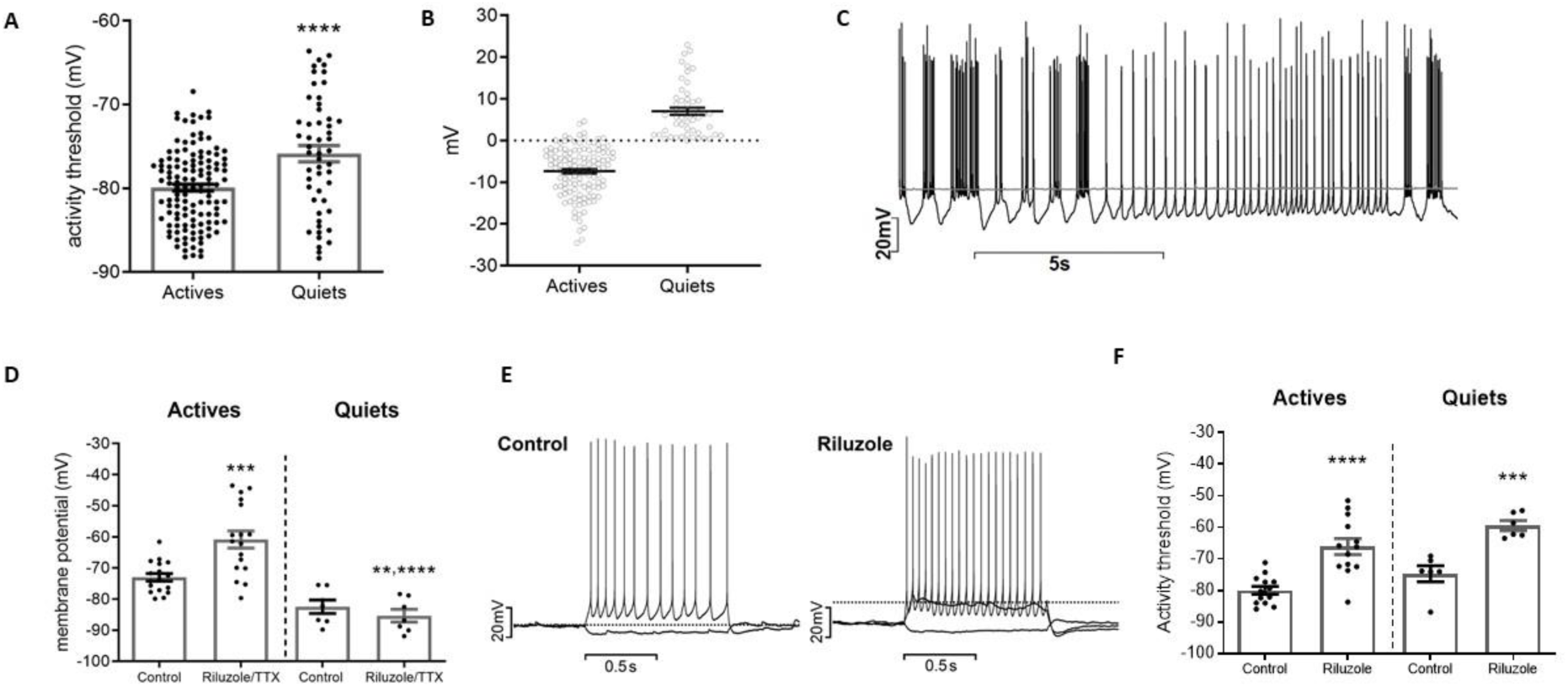
The activity threshold of both quiet and active neurons was affected by the I_NaP_ inhibitor Riluzole. A. activity threshold of cartwheel active and quiet neurons. B. difference between the modal membrane potential (MMP) and activity threshold (AT) in active and quiet neurons (AT - MMP). C membrane potential recording of an active cartwheel neuron before and after (grey trace) application of TTX (0.5 µM). D. modal membrane potential of cartwheel active and quiet neurons before and after application of TTX or riluzole (10 µM). E. evoked firing of a CW neuron before and after the application of riluzole. The dashed line shows the activity threshold. F. activity threshold of active and quiet neurons before and after the application of riluzole. **p<0.01; ***p<0.005; ****p<0.0001.

In active DCN fusiform neurons, riluzole stops spontaneous firing because it raises the activity threshold potential above the resting membrane potential (Leão et al., 2012). To test if I_NaP_ is responsible for the activity threshold in CW neurons, we compared the parameter after riluzole application. Riluzole markedly increased the activity threshold in both active and quiet neurons (Quiet: from −74.8 ± 2.6mV to −59.5 ± 1.6mV, n = 6, p=0.0009; Active: from −80.0 ± 1.3mV to −66.2 ± 2.5mV, n=13, p<0.0001; Figure 3E,F). The activity threshold of quiet and active neurons after riluzole was not significantly different (p=0.1). We conclude that I_NaP_ is responsible for setting the activity threshold of both active and quiet neurons in more hyperpolarized potentials.

We hypothesized that differential expression or changes to the voltage dependence of I_NaP_ could determine the quiet and active mode of the CW neurons, with active neurons expressing more I_NaP_ than quiet neurons. To test this hypothesis, we measured the current sensitive to both riluzole or TTX in quiet and active CW neurons using voltage ramps (Figure 4A, B). Riluzole or TTX abolished the inward component seen in active neurons, allowing the IV to cross the zero current point (Figure 4A). The IV relationships of I_NaP_ was similar in quiet and active neurons (Figure 4B,C), with a current at −50 mV of −433 ± 46pA in quiet neurons and −354 ± 45pA in active neurons (n= 7 and 10 respectively, p=0.25). The I_NaP_ chord conductance (measured at −45 mV) was of 4.2 ± 0.8 nS for quiet neurons and 3.8 ± 0.3nS for active neurons (p=0.48). In IV relationships performed with square pulses, the I_NaP_ has similar parameters as to when measured with the voltage ramps (not shown). Interestingly, the point of zero current of active neurons in the presence of riluzole or TTX was more positive than the point of zero current of quiet neurons (Figure 4A), in accordance to the more depolarized resting membrane potential of active neurons (Figure 3D) and as observed for DCN fusiform neurons (Leão et al., 2012). We conclude that, like DCN fusiform neurons, the expression of I_NaP_ in active and quiet CW neurons is similar, and therefore some other subthreshold conductance must be responsible for determining the quiet and active modes.

**Figure 4:**
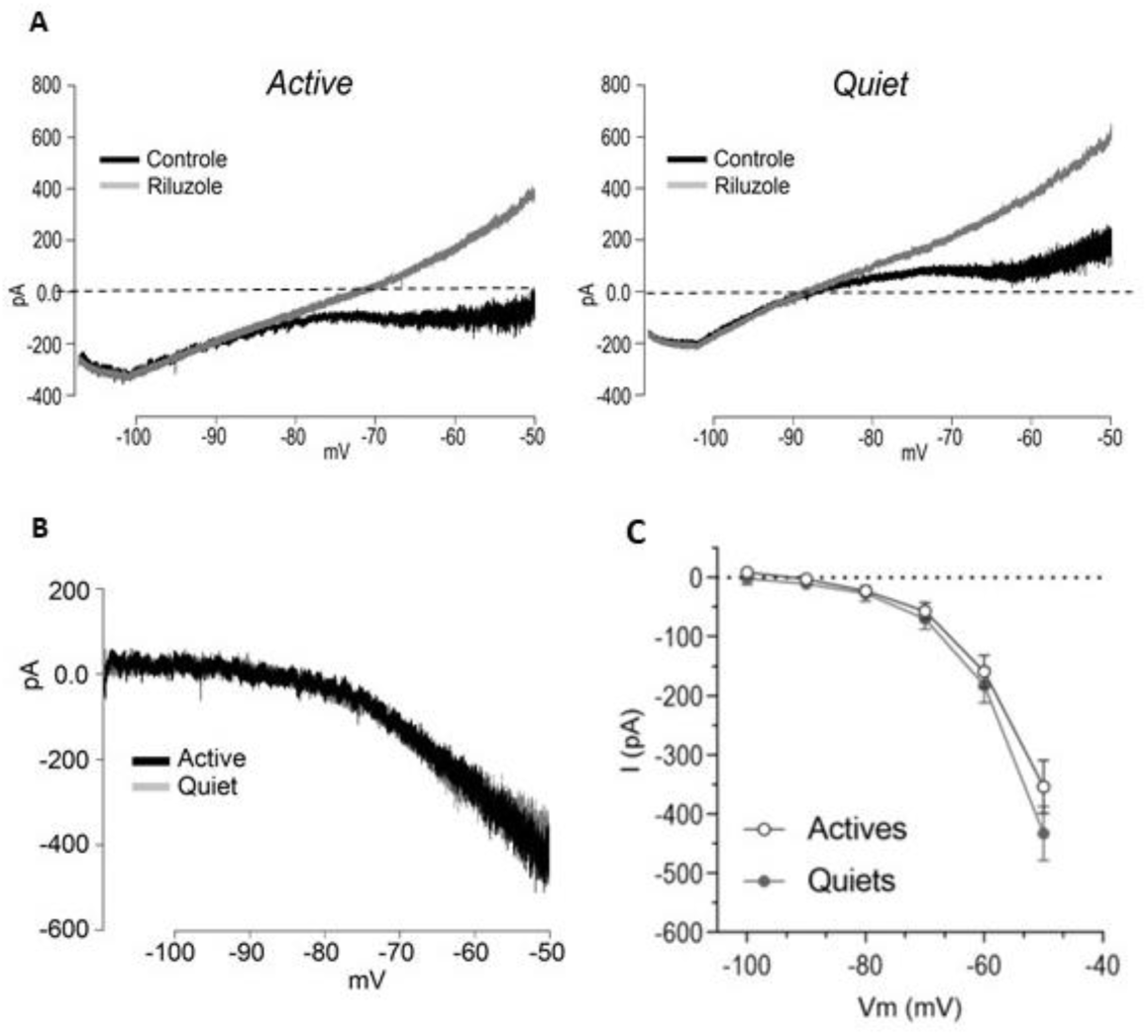
Cartwheel active and quiet neuron equally expresses the persistent sodium current. A. current evoked by ramp commands before and after (grey traces) riluzole (10 µM), of an active and quiet neuron. B. riluzole-sensitive current from an active and quiet (grey trace) neuron. C. IV relationships of sodium persistent currents from quiet and active neurons.

### I_h_ depolarizes the membrane potential of both quiet and active neurons

DCN fusiform neurons express the hyperpolarization-activated cationic current (I_h_) that depolarizes the membrane of both quiet and active neurons, but with no deterministic role in defining these phenotypes (Ceballos et al., 2016). In CW neurons, the inhibition of I_h_ leads to hyperpolarization (Kim and Trussell, 2007). However, the specific role of this current in determining the silent or spontaneous firing neuronal types was not investigated. To test the role of I_h_ in the generation of active and quiet CW neurons, we applied the selective antagonist ZD7288 (20 μM).

Application of ZD7288 hyperpolarized the membrane potential of both quiet and active neurons (Active: from −75.0 ± 1.4 mV to −92.6 ± 0.8 mV, n=24, p<0.0001; Quiet: from −86.7 ± 1.4 mV to −94.7 ± 1.0 mV; n=11, p<0.0001; Figure 5A,B,C). In the presence of ZD, active neurons stopped firing action potential spontaneously. The effect of ZD on membrane potential was stronger for active neurons (−17.6 ± 1.0 mV) than quiet neurons (−8.1 ± 1.0 mV; p<0.0001), and the membrane potential after ZD was similar in both types (p=0.14). As expected, ZD7288 abolished the membrane depolarization sag (characteristic of I_h_) during a hyperpolarization pulse and increased membrane input resistance (Quiet: from 84,7 ± 5.0 MΩ to 126,3± 7,6MΩ, n=12, p<0.0001; Active: from 103.1 ± 3.5 MΩ to 134.2 ± 4.5 MΩ, n=14, p<0.0001; Figure 5D). Interestingly, despite the inhibitory effect on spontaneous firing, ZD7288 decreased the activity threshold of both active and quiet neurons (Quiet: from −80.3 ± 2.0 mV; to −88.0 ± 0.8 mV, n=11, p=0.0004; Active: −82.1 ± 0.8 mV to −87.0 ± 0.8 mV, n=25, p<0.0001; Figure 5E), possibly as a consequence of the increased input resistance. Thus, our results show that I_h_ depolarizes the membrane of both quiet and active CW neurons, and in active neurons, this depolarization crosses the activity threshold leading to spontaneous firing.

**Figure 5:**
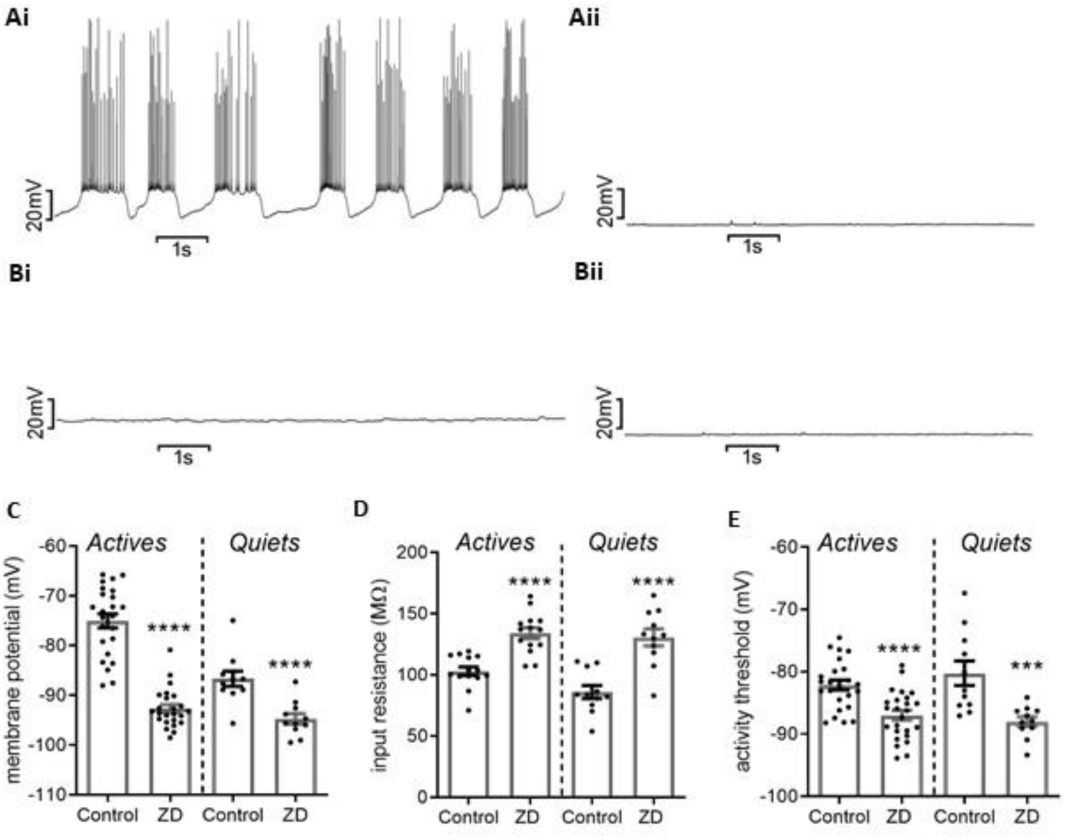
Inhibition of I_h_ hyperpolarizes cartwheel neurons. A. membrane potential recordings of an active cartwheel neuron before (i) and after (ii) application of ZD7288 (20 µM). B. membrane potential recordings of a quiet cartwheel neuron before (i) and after (ii) application of ZD7288. C. membrane potential of active and quiet cartwheel neurons before and after the application of ZD7288. D. membrane input resistance of active and quiet cartwheel neurons before and after the application of ZD7288. E. activity threshold of active and quiet cartwheel neurons before and after the application of ZD7288. ***p<0.005; ****p<0.0001.

Because the effect of ZD7288 was stronger on active neurons, we asked if I_h_ was larger in these neurons than in quiet neurons. We then measured I_h_ in voltage-clamp as the ZD7288-sensitive current (Figure 6A,B). We found that I_h_ was consistently larger in active neurons at all potentials tested (Figure 6C). Accordingly, the chord conductance was significantly bigger in active neurons (Active: 4.1 ± 0.3nS; n=21; Quiet: 3.2 ± 0.3nS; n=9; p=0.036). Interestingly, the I_h_ of quiet neurons is shifted to the left, with the slope conductances not significantly different between groups (Active: 10 ± 0.9 nS; n=9; Quiet: 9.8 ± 0.8 nS; n=8; p=0.91), suggesting a difference in the activation of I_h_ between quiet and active neurons. We conclude that h current is more prominent in active neurons

**Figure 6:**
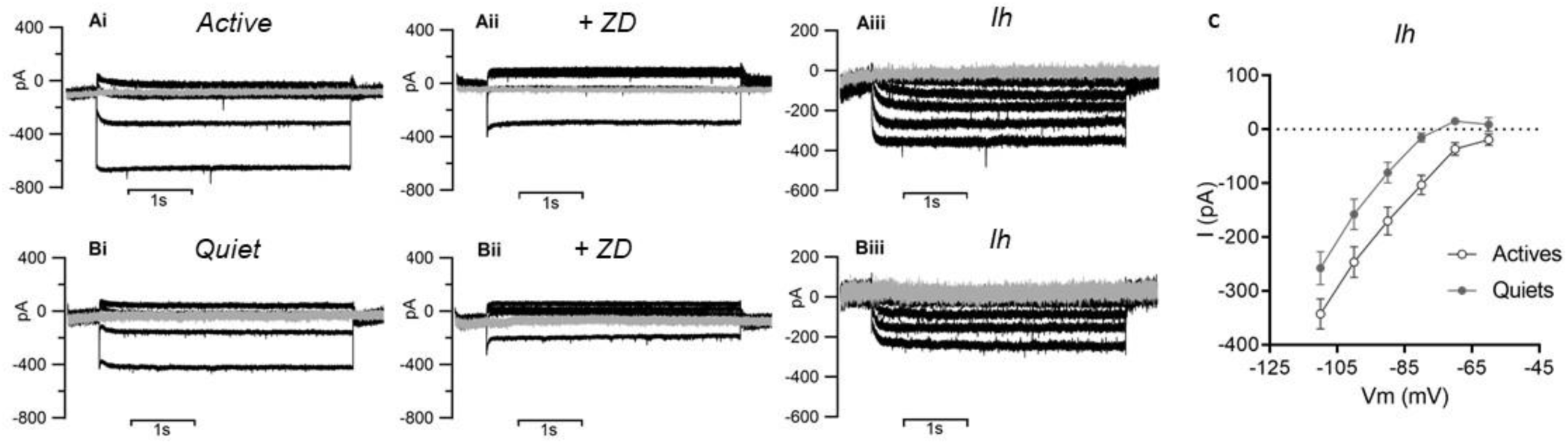
I_h_ is differentially expressed in quiet and active cartwheel neurons. A. membrane subthreshold currents of an active neuron before (i) and after (ii) ZD7288 (20 µM), and the ZD-sensitive current (iii). B. membrane subthreshold currents of a quiet neuron before (i) and after (ii) ZD7288, and the ZD-sensitive current (iii). The grey trace is the holding = −60mV. C. IV relationships of I_h_ of active and quiet cartwheel neurons.

### An inwardly-rectifying potassium current (I_Kir_) controls membrane potential in both active and quiet cartwheel neurons

Looking at the IVs of quiet neurons, it seems that in quiet CW neurons, an outward current counteracts the depolarizing effects of I_NaP_ and I_h_ (Figure 2D), thus avoiding spontaneous firing. In DCN fusiform neurons, the expression of I_Kir_ regulates spontaneous firing, with quiet neurons expressing a more robust I_Kir_ compared to active neurons, which hyperpolarizes the membrane potential below activity threshold produced by I_NaP_ (Leao et al., 2012). To test if CW neurons also make use of I_Kir_ to create active and quiet states, we inhibited I_Kir_ with 100 µM BaCl_2_ (Leao et al., 2012), and compared the Ba^++^-sensitive current in quiet and active neurons.

Perfusion of BaCl_2_ 100 µM strongly depolarized both quiet and active CW neurons (Figure 7D; Quiet: from −81.1 ± 1.7mV to −56.3 ± 4.8mV, n=8, p=0.001; Active: from −70.9 ± 1.4mV to −47.9 ± 3.2mV, n=23, p<0.0001; Figure 7A,B,C,D). The firing behavior after BaCl_2_ 100 µM was complex (Figure 7A,B,C). A fraction of neurons fired action potentials tonically after Ba^++^ (Figure 7A), while another fraction presented bistable behavior (Figure 7B). Interestingly, several neurons depolarized so much that they did not fire action potentials because of depolarization block, by inactivation of sodium channels (Figure 7C). Barium increased the membrane input resistance of active and quiet neurons (Quiet: from 91.7 ± 4.9MΩ to 165.6 ± 14.7MΩ, n=8; p=0,0005; Active: from 114.7 ± 5.1MΩ to 166.7 ± 9.1MΩ, n=20, p<0.0001; Figure 7E), and it became similar in both types (p=0.95).

**Figure 7:**
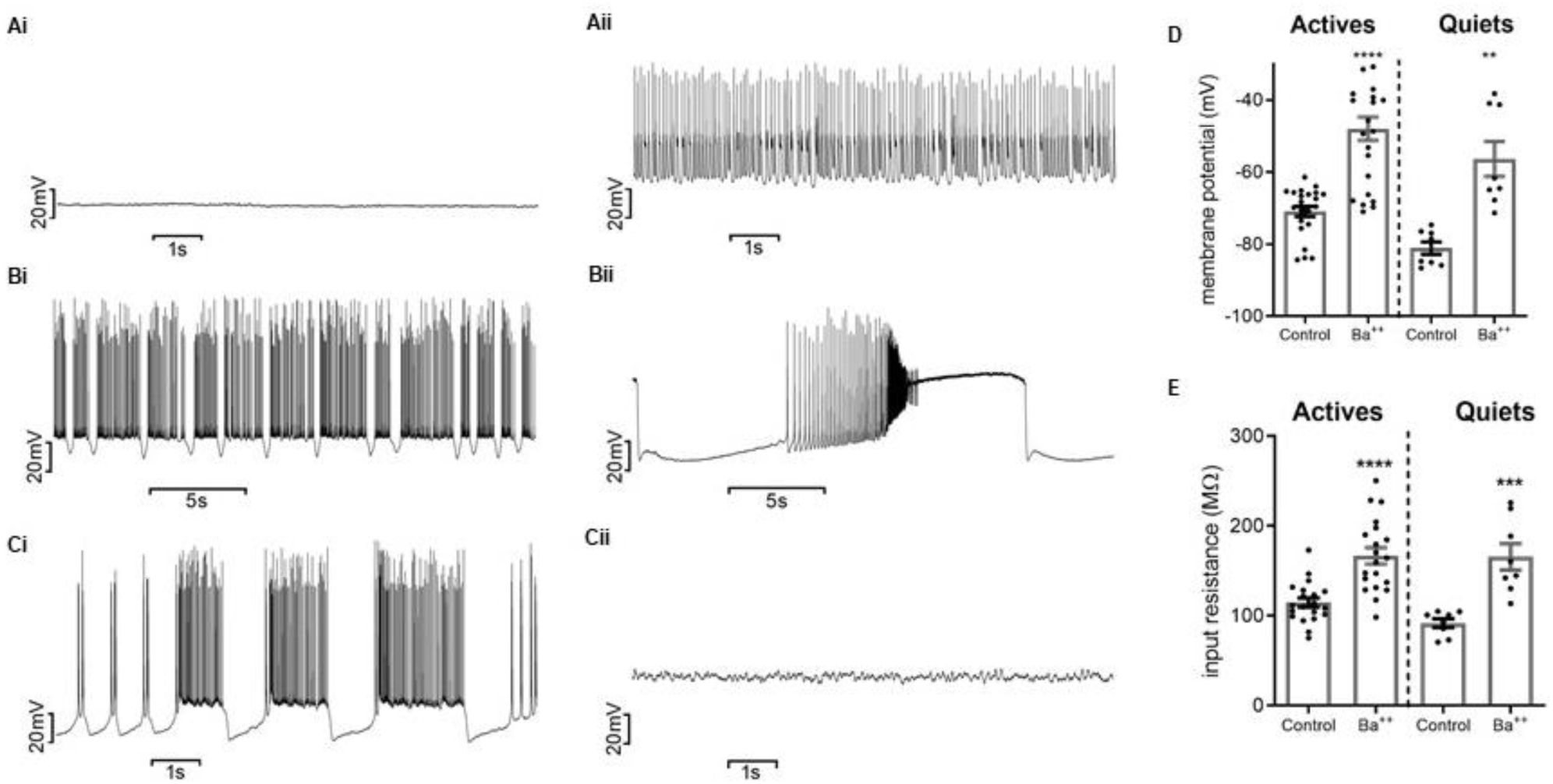
Inhibition of I_Kir_ strongly depolarizes cartwheel neurons. A. membrane potential recordings of a quiet cartwheel neuron before (i) and after (ii) application of BaCl_2_ (0.1 mM). B,C. membrane potential recordings of two active cartwheel neurons before (i) and after (ii) application of BaCl_2_. Bistable behavior is observed in B and depolarization block is observed in C. D. membrane potential of active and quiet cartwheel neurons before and after the application of BaCl_2_. E. membrane input resistance of active and quiet cartwheel neurons before and after the application of BaCl_2_. **p<0.01; ***p<0.005; ****p<0.0001.

To see if there was a differential expression of IK_ir_ in quiet and active CW neurons, we analyzed the membrane currents before and after BaCl_2_ 100 µM in voltage-clamp (Figure 8A,B). We found that barium inhibited the outward component of the IV relationship of quiet CW cells, revealing the inward current of I_NaP_ (Figure 8C). On the other hand, it had a less pronounced effect on active CW neurons, particularly in the more positive potentials (Figure 8D).

**Figure 8:**
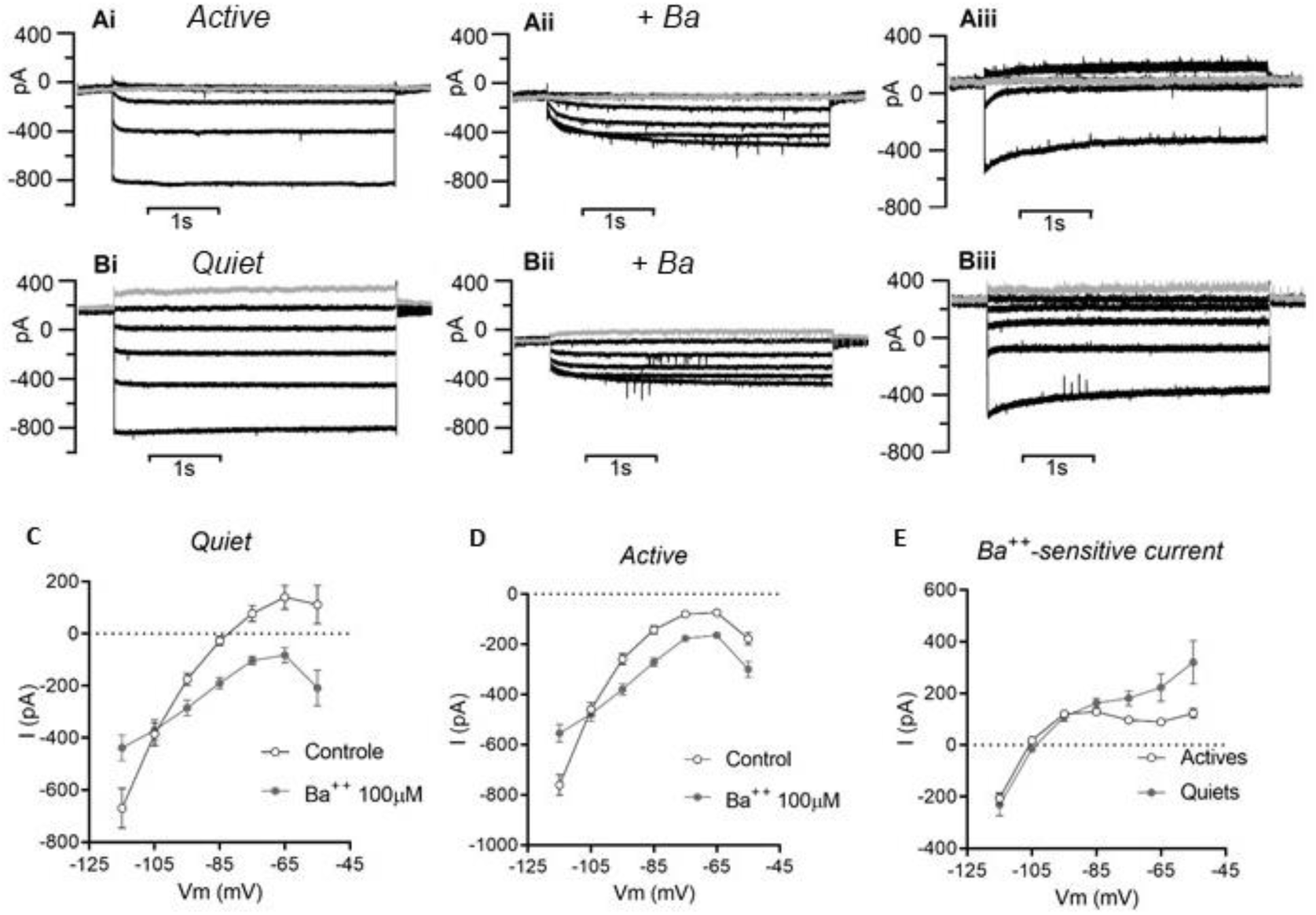
I_Kir_ has different rectifications in quiet and active cartwheel neurons. A. membrane subthreshold currents of an active neuron before (i) and after (ii) BaCl_2_ (0.1 mM), and the barium-sensitive current (iii). B. membrane subthreshold currents of a quiet neuron before (i) and after (ii) BaCl_2_, and the barium-sensitive current (iii). Grey trace is the −65mV step pulse. C. IV relationships before and after BaCl_2_ of quiet cartwheel neurons D. IV relationships before and after BaCl_2_ of active cartwheel neurons. E. IV relationship of I_Kir_ of active and quiet cartwheel neurons.

The barium-sensitive current in active and quiet CW neurons have different voltage dependence, as can be seen in figure 8E. While in active neurons, the barium sensitive current has the profile of a classic I_Kir_, with strong rectification, in quiet neurons, the current has less rectification with a robust outward component (Figure 8E). On the other hand, in the range of −115 to −95 mV, the current was similar in both quiet and active CW cells (Figure 8E). The slope conductance measured in this region was similar in both types (quiet: 17.2 ± 2.5 nS; Active: 15.8 ± 1.5 nS; n=6 and 20, respectively; p=0.7). We conclude that in both quiet and active neurons, I_Kir_ strongly regulates membrane potential. However, quiet neurons express an I_Kir_ with a weak inward rectification that counteracts the inward currents of I_NaP_ and I_h_, preventing the membrane potential from crossing the activity threshold and, thus, avoiding spontaneous action potential firing.

### An outward current from ATP-sensitive potassium channels (K_ATP_) is present in quiet cartwheel neurons and prevents spontaneous firing

We observed that barium-sensitive currents in quiet CW neurons present a more pronounced outward current near the activity threshold than in active neurons. This result could reflect a smaller rectification of I_Kir_ currents in quiet neurons, or some other low-threshold K_v_ current sensitive to BaCl_2_ 100 µM. K_ATP_ channels belong to the family of inward rectifying potassium channels, and its currents present less rectification than other types of K_ir_ channels (Bichet et al., 2003). K_ATP_ currents are produced by the subunits K_ir_6.2 and 6.1, which are expressed in several central regions (Dunn-Meynell, 1998; Zhou et al., 1999, 2002). We, therefore, tested if the difference in the rectification of the Ba^++^ 100 µM-sensitive current of quiet and active CW neurons was caused by differential expression of K_ATP_ currents.

To test the presence of K_ATP_ currents in CW neurons, we perfused the K_ATP_ channel antagonist tolbutamide (100 µM) on quiet and active CW neurons. The application of tolbutamide did not change the behavior of most active neurons (Figure 9A). Only one out of 16 neurons noticeably increased action potential firing rate and membrane input resistance after tolbutamide. In the other 15 neurons, tolbutamide did not significantly change membrane potential (from −70.6 ± 1.0mV to: −69.5 ± 1.4mV, n=14, p=0.21; Figure 9C) and input resistance (from 115.3 ± 5.6MΩ to 114.2 ± 5.3 MΩ; n=15, p=0,68; Figure 9D), and only slightly reduced activity threshold (from 77.6 ± 1.1mV to −79 ± 1.1mV; n=14, p=0,055).

**Figure 9:**
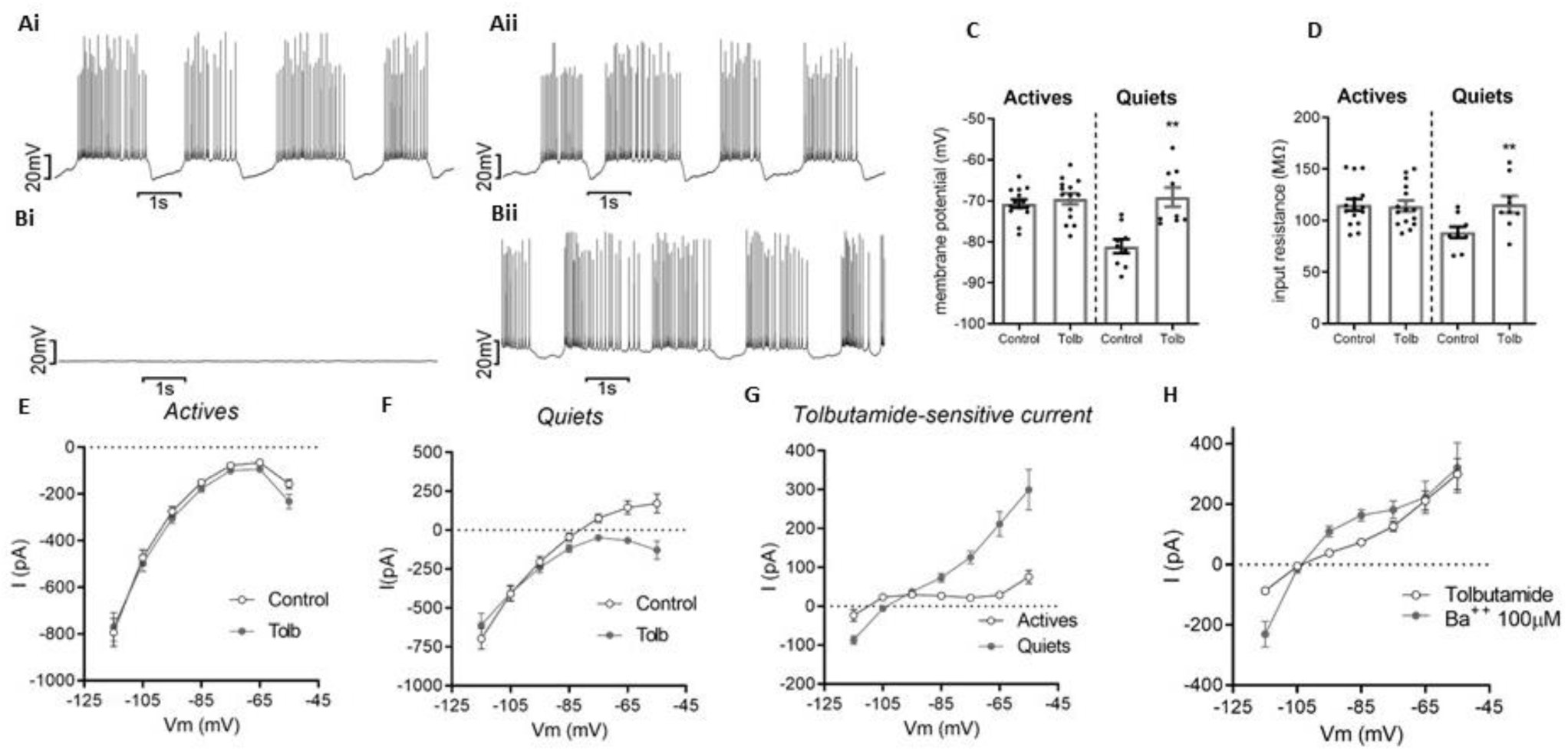
Quiet cartwheel neurons express open K_ATP_ channels. A. membrane potential recordings of an active cartwheel neuron before (i) and after (ii) application tolbutamide (0.1mM). B. membrane potential recordings of a quiet cartwheel neuron before (i) and after (ii) application of tolbutamide. C. membrane potential of active and quiet cartwheel neurons before and after the application of tolbutamide. D. membrane input resistance of active and quiet cartwheel neurons before and after the application of tolbutamide. E. IV relationships before and after tolbutamide of active cartwheel neurons. F. IV relationships before and after tolbutamide of quiet cartwheel neurons. G. IV relationship of tolbutamide-sensitive current of active and quiet cartwheel neurons. H. Comparison of the tolbutamide-sensitive current and the BaCl_2_ (0.1 mM) sensitive current in quiet cartwheel neurons. **p<0.01.

On the other hand, most quiet neurons (9 out of 11, or 82%) started to fire spontaneously after tolbutamide (Figure 9B), most of them with firing patterns similar to that of control active neurons. Tolbutamide depolarized quiet neurons from −81.1 ± 1,7mV to −69.1 ± 2.3mV (n=9, p=0.0035; Figure 9C) and increased input resistance from 88.7 ± 5.4 MΩ to 115.8 ± 8.2 MΩ (n=9. p=0.0015; Figure 9D). It also decreased the activity threshold from −73.3 ± 2.0 mV to −80.6 ± 1.2mV (n=9, p=0.0023). We did not find significant differences in any of these parameters when comparing the quiet group after tolbutamide with the control active group.

We then analyzed the ionic currents affected by tolbutamide in voltage-clamp. While tolbutamide did not affect the membrane currents of active CW neurons (Figure 9E), it inhibited the outward component of the IV of quiet neurons, revealing the developing of the inward I_NaP_ before the point of zero current (Figure 9F), as seen in the IVs of control active neurons. The tolbutamide-sensitive current has little to no rectification, like the current inhibited by BaCl_2_ 100μM in quiet neurons (figure 9G). Comparing the currents sensitive to barium and tolbutamide in quiet neurons, we found that they had similar magnitudes at the range of −65 and −55 mV. However, the tolbutamide-sensitive current had a smaller magnitude in the voltage range from −115 mV to −75mV (Figure 9H). We conclude that part of the potassium inwardly-rectifying current in quiet neurons is composed of open K_ATP_ channels and that the IK_ATP_ is responsible for maintaining the resting membrane potential by counteracting the depolarizing drive of the persistent sodium current.

### K_ATP_ channels are present in active cartwheel neurons and IK_ATP_ suppresses spontaneous firing

Our next question was if only quiet CW neurons express K_ATP_ channels or if these channels are present but closed in active CW neurons. For this, we applied an agonist of K_ATP_ channels, diazoxide, which opens K_ATP_ channels even when they are blocked by intracellular ATP (Nichols, 2006). Application of diazoxide (200 µM) hyperpolarized the membrane in 15 of 17 active CW neurons, eliminating (n=13) or reducing (n=2) their spontaneous firing (Figure 10A). Diazoxide hyperpolarized the membrane potential from −69.8 ± 1.5mV to −81.3 ± 1.6 mV (n=15; p<0.0001; Figure 10B), reduced the membrane input resistance (from 113.5 ± 3.8 MΩ to 80.3 ± 3.6MΩ; n=15, p<0.0001; Figure 10C) and increased the activity threshold (from −81.3 ± 1.0mV to −72.4 ± 1.6mV; n=16, p<0.0001). Interestingly, these parameters are not significantly different in comparison to quiet neurons (p>0.05; figure 10B, C).

**Figure 10:**
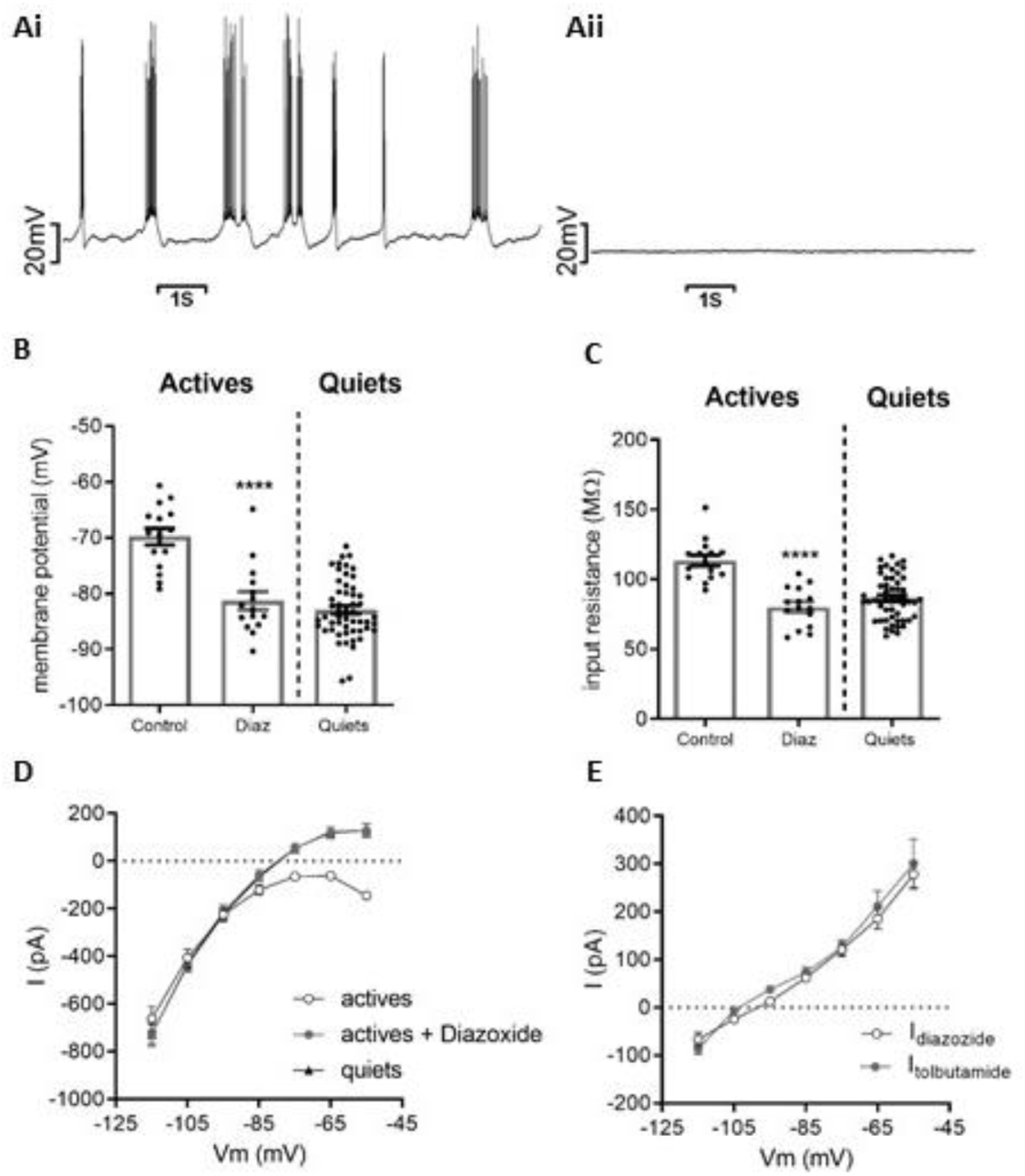
Active cartwheel neurons express closed K_ATP_ channels. A. membrane potential recordings of an active cartwheel neuron before (i) and after (ii) application of diazoxide (0.2 mM). B. membrane potential of active cartwheel neurons before and after application of diazoxide and of control quiet cartwheel neurons. C. membrane input resistance of active cartwheel neurons before and after application of diazoxide and of control quiet cartwheel neurons. D. IV relationships before and after diazoxide of active cartwheel neurons, and comparison with the IV of control quiet neurons. E. Comparison of the diazoxide-sensitive current in active cartwheel neurons with the tolbutamide-sensitive current in quiet neurons. ****p<0.0001.

We then performed voltage-clamp recordings of membrane currents after diazoxide. We found that the application of diazoxide to active neurons changed the IV relationship to a shape almost identical to the IV of control quiet neurons (Figure 10D). Strikingly, we found that the diazoxide-activated current of active neurons was similar in shape and magnitude to the tolbutamide-sensitive current of quiet neurons (Figure 10E), and the chord conductances of these currents were not significantly different (Quiet: 5.5 ± 0.9 nS, n=8; Active: 5.3 ± 0.5 nS, n=15; p=0.9). We conclude that K_ATP_ channels are present in both quiet and active CW neurons but are open in quiet neurons and closed in active neurons, and this difference is responsible for the quiet/active CW neuronal firing modes.

### Glycinergic neurotransmission on the fusiform neuron is affected by K_ATP_ channel activity

Because most cartwheel neurons fire spontaneously, different from the tuberculoventral neurons, which are silent at rest, they provide a robust glycinergic tone to the fusiform neuron. Thus, if K_ATP_ channels are responsible for creating the quiet mode, we should observe an increase in the glycinergic neurotransmission on the DCN fusiform neuron after the application of tolbutamide and decrease by application of diazoxide. We then applied tolbutamide or diazoxide to DCN slices while recording glycinergic sIPSCs from fusiform neurons (Figure 11A, D). According to the existence of quiet cartwheel neurons, the application of tolbutamide increased the frequency of sIPSCs on fusiform neurons from 17.7 ± 2 Hz to 24.1 ± 3 Hz (n=7; p=0.01. Figure 11B). The amplitude of the sIPSCs was also slightly increased by tolbutamide (from 52.2 ± 7 pA to 61.9 ± 7 pA; n=7. p=0.059. Figure 11C). On the other hand, activation of K_ATP_ channels with diazoxide decreased sIPSCs frequency from 18.6 ± 1.2 Hz to 11.5 ± 1.4 Hz (n=5, p=0.002; Figure 11D) and amplitude from 60.4 ± 6.6 pA to 40.9 ± 2.5 pA (n=5, p=0.02; Figure 11E). We conclude that the control of quiet and active modes of cartwheel neurons by K_ATP_ channels strongly modulates inhibitory neurotransmission on DCN fusiform neurons.

**Figure 11:**
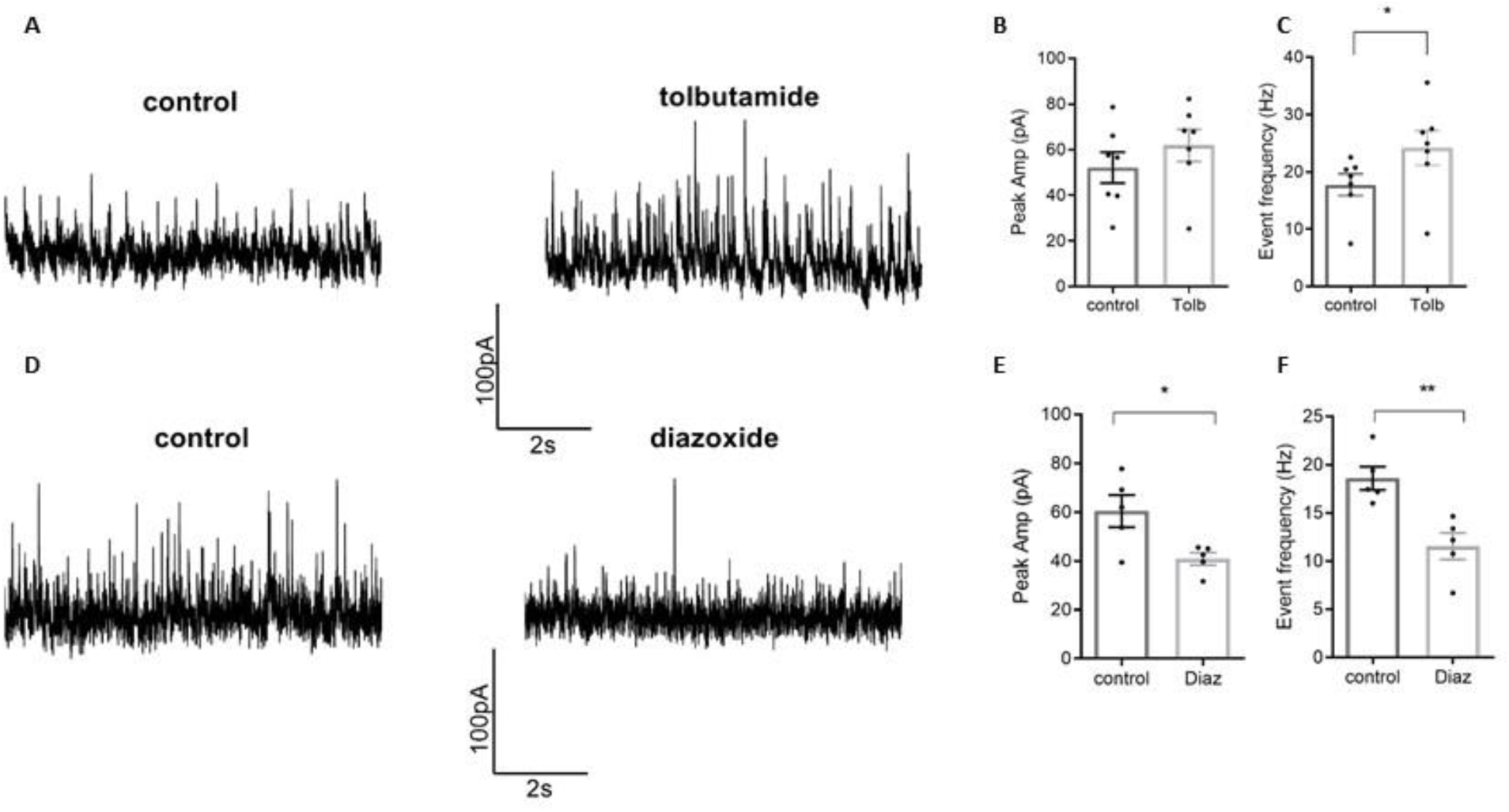
Spontaneous glycinergic neurotransmission on fusiform neurons is modulated by K_ATP_ channels. A. representative traces of sIPSCs in control conditions and after application of tolbutamide (0.1mM). B. sIPSCs mean amplitude before (control) and after application of tolbutamide. C. frequency of sIPSCs before (control) and after application of tolbutamide. D. representative traces of sIPSCs in control conditions and after application of diazoxide. E. sIPSCs mean amplitude before (control) and after application of diazoxide. F. frequency of sIPSCs before (control) and after application of diazoxide. *p<0.05; **p<0.01.

### The proportion of quiet and active cartwheel neurons and the spontaneous glycinergic neurotransmission on fusiform neurons is not affected by extracellular glucose

We performed our recordings using an external solution with 10 mM glucose, which is well above plasmatic normoglycemic values, which lie around 5 mM. *In vitro* experiments in neurons from the nucleus of the solitary tract (NTS) showed that the membrane potential of these neurons is sensitive to external glucose, being more depolarized in 10 mM external glucose than in 5 mM, because of K_ATP_ channel blockade (de Bernardis Murat and Leão, 2019). Thus, we might be overestimating the proportion of cartwheel active neurons than found physiologically, since we used glucose concentrations twice the normoglycemic value. We then performed recordings of cartwheel neurons using aCSF with 5 mM glucose, to see if the proportion of quiet neurons would be bigger in normoglycemic levels of glucose. Contrary to our expectation, the proportion of quiet and active neurons was the same in 5 mM glucose, with 20% of quiet neurons versus 80% of active neurons (n=4 quiet, and n=16 active; p = 0.56; Fisher’s exact test). The membrane potentials of quiet neurons were more hyperpolarized than active neurons, similar to observed in 10 mM glucose (active: −70.4 ± 1.7 mV, quiet: −79.2 ± 4.6 mV, p = 0.048, Figure 12A). The IV relationships were also similar to observed in 10 mM glucose, and the application of tolbutamide change the IV relationship of quiet cells to the profile seen in active cells (Figure 12B), and produced spontaneous firing. We also compared the glycinergic neurotransmission on the fusiform neuron in 5 mM glucose and found that the frequency and amplitude of sIPSCs were similar to in 10 mM glucose (Figure 12C). We conclude that differently to neurons in NTS, the resting membrane potential of cartwheel neurons is not sensitive to variations of external glucose from 5 to 10 mM, and consequently, the quiet/active neuron ratio and spontaneous glycinergic neurotransmission are the same in 5 or 10 mM glucose.

**Figure 12:**
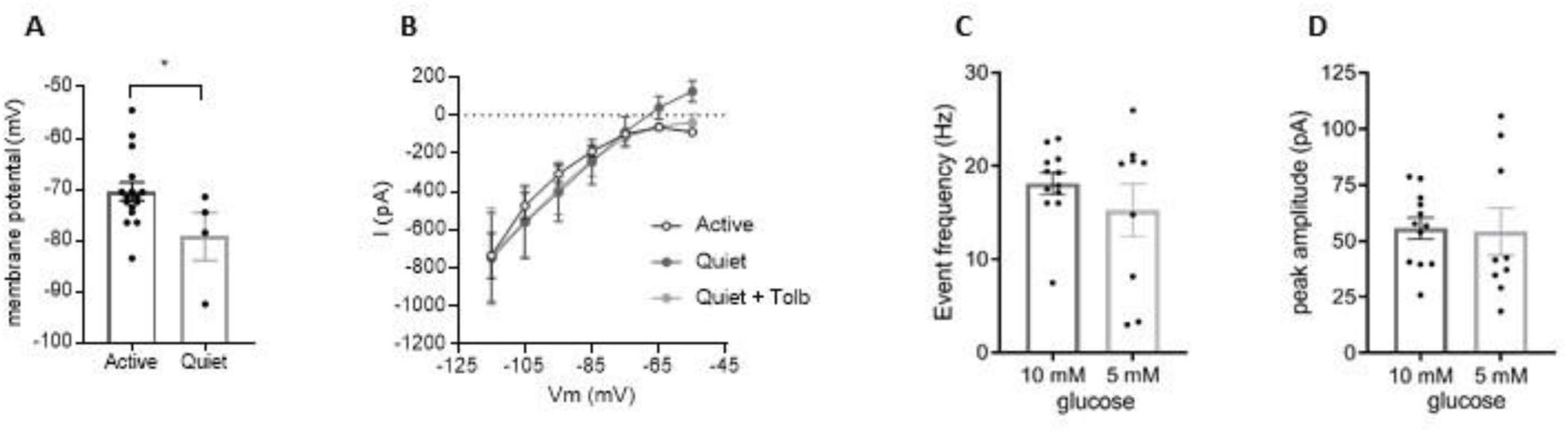
Incubation in 5 mM glucose does not change the membrane potential of cartwheel neurons and glycinergic neurotransmission on fusiform neurons. A. membrane potential of quiet and active cartwheel neurons in 5 mM external glucose. B. IV relationship of quiet and active cartwheel neurons in 5 mM external glucose, and the effect of tolbutamide (0.1mM) on quiet neurons IV. C. sIPSCs frequency in 10 mM and 5 mM external glucose. C. sIPSCs mean amplitude in 10 mM and 5 mM external glucose.

### High doses of salicylate increase the proportion of quiet CW neurons by opening K_ATP_ channels

We previously have shown that incubation of DCN slices with high doses of salicylate increased the proportion of quiet CW neurons (Zugaib et al., 2016). However, the mechanism behind this effect was not investigated. In order to know the ionic mechanisms underlying this effect, we recorded the IV relationships of quiet and active neurons in DCN slices incubated and recorded in the presence of 1.4 mM sodium salicylate as in Zugaib et al. (2016). We again found an increased proportion of quiet CW neurons in the presence of salicylate (quiet: 44%; active 56%; n=50) when compared with control aCSF (18.5% quiets and 81.5% actives; p=0.0004; Figure 13A). The membrane properties of active and quiet neurons in the presence of salicylate were similar to that of control quiet and active neurons (table 1).

**Table 1:** membrane properties of quiet and active CW neurons in control conditions and in the presence of salicylate 1.4 mM. The sample size (n) is in parentheses. The asterisk (*) indicates statistical difference (p < 0.05; unpaired t-test). R_input_= membrane input resistance; MMP= modal membrane potential; AT=activity threshold; tau=membrane time constant; sag= depolarization sag.

|  | Active |  | Quiet |  |
| --- | --- | --- | --- | --- |
|  | Control | Salicylate (n=28) | Control (n=52) | Salicylate (n=22) |
| **R<sub>input</sub> (MΩ)** | 116.3 ± 1.9 (123) | 112.4 ± 3 | 86.5 ± 2.2 | 79.3 ± 3 |
| **MMP (mV)** | -72.5 ± 0.6 (133) | -72.3 ± 1 | -82.9 ± 0.7 | -84.1 ± 1 |
| **AT (mV)** | -79.9 ± 0.4 (124) | -82.8 ± 0.9 * | -75.9 ± 0.9 | -74.8 ± 2 |
| **tau (ms)** | 21.6 ± 0.5 (121) | 20.7 ± 1 | 13.1 ± 0.5 | 11.7 ± 0.7 |
| **sag (mV)** | 2.1 ± 0.1 (120) | 2.2 ± 0.3 | 1.3 ± 0.1 | 1.3 ± 0.1 |

**Figure 13:**
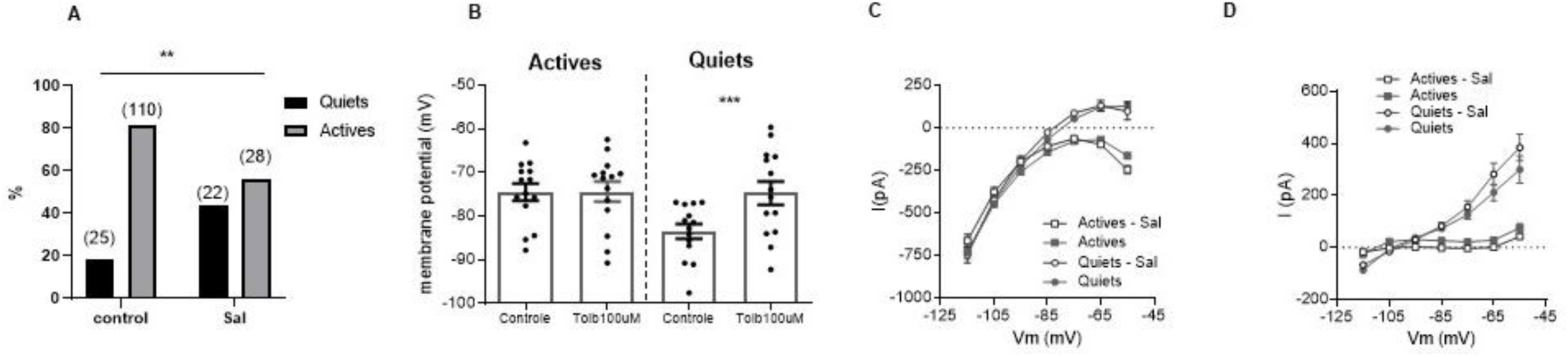
Salicylate increases the number of quiet neurons by opening K_ATP_ channels. A. proportion of quiet and active cartwheel neurons in control aCSF and in the presence of 1.4 mM sodium salicylate (sample size in parentheses). B. membrane potential of active and quiet cartwheel neurons in the presence of 1.4 mM sodium salicylate before and after the application of tolbutamide. C. IV relationships of quiet and active cartwheel neurons in control aCSF and aCSF with 1.4 mM sodium salicylate. D. Tolbutamide-sensitive currents of quiet and active cartwheel neurons in control aCSF and aCSF with 1.4 mM sodium salicylate. **p<0.01; ***p<0.005.

We then tested if the membrane potential of quiet neurons in the presence of salicylate was sensitive to tolbutamide. We found that the application of tolbutamide depolarized the membrane of most quiet CW neurons (14 out of 17, 82%) (Figure 13B), which started firing spontaneously, except by one that depolarized above the AP threshold (causing inactivation of the voltage-dependent Na^+^ channels). In general, the firing patterns acquired were again similar to the ones observed in control active neurons. On the other hand, the active neurons recorded in the presence of salicylate were unaffected by tolbutamide (Figure 13B).

We then examined the IV relationships of active and quiet neurons in the presence of salicylate. The IV profile of quiet and active neurons in salicylate was very similar to the IV of control aCSF neurons (Figure 13 C). Additionally, the effect of tolbutamide in the IV curves was the same on quiet neurons in salicylate, with no impact in active neurons (not shown). The tolbutamide-sensitive currents were similar in the control aCSF and salicylate group (Figure 13D). We then conclude that high doses of salicylate increase the proportion of quiet CW neurons by increasing the opening of K_ATP_ channels.

## Discussion

In this report, we described a new role of K_ATP_ channels in the control of neuronal firing mode. Our motivation for this research was to investigate the ionic mechanisms underlying the quiet and active states of CW neurons from the DCN. Because of its central role in the inhibition of the fusiform neuron, knowing the ionic channels controlling the firing of the glycinergic CW cell is central to understand the mechanisms of the hyperexcitability observed in the fusiform neuron in animal models of tinnitus (Kaltenbach and Afman., 2000; Brozoski et al., 2002), which can be related to a decrease in the glycinergic inhibition from CW neurons (Zugaib et al., 2016, Zugaib and Leão, 2018).

Previous reports showed that most CW neurons fire spontaneous action potentials (Kim and Trussell, 2007; Zugaib et al., 2016). Kim and Trussell (2007), using perforate-patch recordings from mice CW neurons, found 66% of active neurons and 34% of quiet neurons. We found the proportion of 81.5% active cells and 18.5% of quiet cells. We thus found a smaller proportion of quiet neurons than the previous reports, and contingency analysis of our data with the data from Kim and Trussell (2007) showed a significant difference in the proportions (p=0.001; Fisher’s exact test). This difference could be related to the species (rat versus mice) and the method employed (perforated patch versus whole-cell). Using rats and whole-cell patch-clamp, Zugaib et al. (2016) found that 73% of neurons were active and 27% quiet, an intermediate proportion. We can conclude that despite these differences, CW neurons are mostly active in both mice and rats and because we found a similar proportion of active and quiet neurons in cell-attached experiments, the prevalence of the active state in CW neurons does not seem to be caused by artifacts from the whole-cell configuration.

Fusiform neurons from the DCN present both quiet and active modes, but in equal proportions (Leão et al., 2012; Zugaib et al., 2016). In fusiform neurons, the spontaneous firing is driven by the I_NaP_, but its expression is similar in quiet and active neurons. We found similar results in CW neurons showing that, like in fusiform neurons, there is no differential expression of I_NaP_ that could explain the two firing modes. Interestingly, inhibition of AP firing with riluzole or TTX depolarized the modal membrane potential of active neurons, probably by decreasing calcium entry during action potential firing, thus inhibiting calcium-activated potassium channels, which produces strong post-potential hyperpolarizations (Kim and Trussel, 2007).

Kim and Trussell (2007) suggested that a balance between the hyperpolarization-activated cationic current (I_h_) and the inward rectifying potassium current (I_kir_) could be important in determining the level of activity of CW neurons. Our previous results with fusiform neurons showed an increased expression of the h current in active neurons, but with no deterministic role in generating the quiet/active modes of fusiform neurons (Ceballos et al., 2016). We similarly found an increased I_h_ in active CW neurons. This difference could be related to differences in the modulatory state of the HCN channels, since we observed a shift to the left of I_h_ IV of quiet neurons, suggestive to the effect of cAMP on the HCN channels (Biel et al., 2009). Inhibition of I_h_ stopped the spontaneous firing of active neurons, showing that the basal depolarization produced by I_h_ is essential for generating spontaneous firing. These results could lead to the conclusion that the differential expression of I_h_ in active and quiet neurons would lead to the creation of the two modes. However, the presence of a robust outward component in the IV curves of quiet neurons suggested that another conductance would play a role in curtailing the spontaneous firing of these neurons.

Because our previous results in fusiform neurons indicated that I_Kir_ expression was more prominent in quiet neurons and responsible for creating the quiet mode (Leão et al., 2012), we inhibited I_Kir_ in CW neurons with 100 μM barium and found that both quiet and active neurons strongly depolarized after barium application. However, the analysis of the barium-sensitive current revealed distinct voltage dependencies of the current recorded in quiet versus active neurons. The barium-sensitive current in quiet neurons showed an outward component, starting from −85mV, which was much smaller in active neurons (Figure 8E), suggesting that a potassium channel sensitive to micromolar concentrations, of barium with no or little rectification is present in quiet neurons. As the first hypotheses, we tested for IK_ATP_, a current belonging to the family of inward rectifying potassium currents and known to present weaker inward rectification (Bichet et al., 2003). K_ir_6.1/2 subunits express KATP channels along with the auxiliary SUR subunits, which are necessary for their sensitivity to sulphonylureas, as tolbutamide (Bichet et al., 2003). Based on their sensitivity to tolbutamide and diazoxide the CW neurons probably express Kir6.2/SUR1 subunits (Grible et al., 1998; Hansen, 2006; Friedland and Popper, 2007).

We found that only quiet CW neurons expressed a tolbutamide-sensitive current, and they became active in the presence of tolbutamide. Interestingly, the effect of tolbutamide on firing was not as dramatic as the effect of barium, with most quiet neurons changing to firing patterns and frequency similar to observed in control active neurons, in accordance with the presence of other Kir channels controlling membrane potential. However, in a few quiet neurons, we did not observe the effect of tolbutamide to depolarize the membrane potential, suggesting that in a few CW neurons K_ATP_ is not pivotal to produce the quiet mode. The activation of IK_ATP_ with similar IV dependence in active neurons by diazoxide showed that active CW neurons express K_ATP_ channels but in the closed state. Strikingly, activation of these channels by diazoxide turned them to quiet, producing an IV relationship identical to that of control quiet neurons. Interestingly, very few active neurons were unresponsive to diazoxide or maintained spontaneous firing after the drug application, showing that there are secondary mechanisms that can contribute to the generation of the active and quiet CW neurons.

K_ATP_ channels are blocked by ATP and activated by ADP, which allows them to serve as sensors of the energetic state of the cell (Nichols, 2006). The membrane lipid PIP_2_ (Phosphatidylinositol 4,5-bisphosphate) decreases their sensitivity to ATP (Baukrowitz et al., 1998; Shyng et al., 2000), making these channels open in millimolar concentrations of intracellular ATP. We used whole-cell patch-clamp with millimolar concentrations (2 mM) of ATP in the internal solution, which could mask the physiological levels of ATP. Thus, it is surprising that we found a fraction of cells with open K_ATP_ channels. However, other reports have shown that K_ATP_ channels are more sensitive to metabolic ATP than to the ATP supplied by the electrode (Song et al., 2001; Muller et al., 2002; Balfour et al., 2006; De Bernardis Murat and Leão, 2019), probably by compartmentalization of K_ATP_ channels and the ATP-producing enzymes (Garg et al., 2009; Zecchin et al., 2015). However, it is conceivable that there is a more significant proportion of cells with closed K_ATP_ channels by the ATP provided by the electrode. Nonetheless, as mentioned before, we obtained the same proportion of active and quiet CW neurons using the cell-attached technique. Thus the ATP in the electrode may have little or no influence in our whole-cell experimental results.

We found that K_ATP_ activity was not sensitive to decreasing external glucose from the high levels traditionally used in slice recordings (10 mM) to concentrations closer to normoglycemic levels (5 mM). We recently demonstrated that neurons of the nucleus of the solitary tract (NTS) are more depolarized when incubated in 10 mM of glucose than in 5 mM of glucose because of more K_ATP_ channels blocked in the high glucose solution (de Bernardis Murat, 2019), showing that these neurons are sensitive to metabolic ATP, even when tested in whole-cell patch-clamp with ATP in the internal solution. However, when we incubated DCN slices in 5 mM glucose, we did not observe differences in the relationship of quiet and active CW neurons, and the spontaneous glycinergic neurotransmission in the fusiform neuron, compared to neurons in 10 mM glucose. These findings suggest that cartwheel neurons may not be as sensitive as NTS neurons to possible changes in metabolic ATP caused by changes in external glucose. Nevertheless, more careful experiments using perforated patch-clamp are necessary to understand the role of metabolic ATP in controlling the membrane potential of cartwheel neurons.

We found that two conductances are differentially expressed in CW active and quiet neurons, the I_h_ and the IK_ATP._ I_h_ inhibition prevents active neurons from firing spontaneously, and K_ATP_ inhibition turns quiet neurons into active neurons, while K_ATP_ activation turns active neurons into quiet. Which conductance, I_h_, or K_ATP_, is more critical to determine the quiet/active phenotype? The combination of both conductances is necessary to produce quiet and active CW neurons. Closing of K_ATP_ channels favors the active phenotype, whereas activation of K_ATP_ favors the quiet phenotype. Activation of K_ATP_ channels by diazoxide produces a current in active neurons very similar to IK_ATP_ in quiet neurons, which is enough to stop the firing of most of active neurons, despite their larger I_h_. Additionally, even though quiet neurons have a smaller I_h_, it is sufficient to promote spontaneous firing when IK_ATP_ is inhibited by tolbutamide. It is possible, however, that in the small fraction of quiet neurons unresponsive to tolbutamide, I_h_ is an important factor contributing to the quiet state. Thus, we conclude that open K_ATP_ channels is the principal determinant of the quiet state in CW neurons, and closed K_ATP_ channels, of the active state. This finding is surprising because K_ATP_ channels are not usually associated with control of firing in central neurons. We do not know yet if K_ATP_ channels activation is essential for signal processing in the DCN or a consequence of ATP depletion after intense firing, which could be a way of reducing the energy demands needed to maintain the ionic gradients during prolonged action potential firing. More studies on the sensitivity of the K_ATP_ currents of CW neurons to the metabolic status would be necessary to know if their activity relates to the energetic demands of the neuron.

CW neurons provide a strong inhibitory drive to fusiform neurons and knowing which channels control the firing of these neurons is relevant for understanding the control of the excitability of fusiform neurons, whose hyperactivity is associated with tinnitus (Kaltenbach and Afman, 2000; Brozoski et al., 2002). Consistent to what we have found in CW neurons, the spontaneous glycinergic neurotransmission on fusiform neurons (which is mainly driven by CW neurons spontaneous firing) was sensitive to tolbutamide and diazoxide, showing that both open and closed K_ATP_ channels are present in intact CW neurons and influence their firing. These results also show that the regulation of K_ATP_ channel influences the inhibitory drive on the fusiform neuron. Interestingly, we found a similar magnitude of the effects of tolbutamide and diazoxide on the sIPSC frequency (tolbutamide: 6.4 ± 2 Hz; diazoxide: - 7.1 + 0.9 Hz), which was not expected, because that the existence of a bigger proportion of quiet neuron would result in a stronger inhibitory effect of diazoxide than a stimulatory effect of tolbutamide. At first glance, this could be interpreted that the real proportion of quiet neurons is bigger than we measured. However, it must be taken into account the synaptic contacts of cartwheel neurons, which make synapses with themselves, which have evidence to be excitatory (Tzounopoulos et al., 2004), which could lead to non-linear effects of tolbutamide and diazoxide in the sIPSCs frequency.

Sodium salicylate in high doses induces tinnitus (Stolzberg et al., 2012). When incubated in slices of DCN, salicylate greatly increased the proportion of quiet CW neurons (from 27% to 62%) and simultaneously reduced spontaneous glycinergic neurotransmission on fusiform neurons from 19.0 to 11.2Hz (Zugaib et al., 2016), an effect similar to that produced by diazoxide in our experiments. However, the mechanisms by which salicylate increased the proportion of quiet neurons were unknown. We found that treatment with salicylate activates K_ATP_ channels, thus increasing the fraction of quiet neurons. We do not know the mechanisms by which salicylate produces this effect. In high doses, salicylate is a mitochondrial uncoupler (Haas et al., 1985), which could lead to a decrease in cytoplasmic ATP levels near K_ATP_ channels. Additionally, salicylate in millimolar concentrations is an activator of the AMP-activated protein kinase (AMPK; Hawley et al., 2012), which can activate K_ATP_ channels (Beall et al., 2003; Wu et al., 2017). Thus, it is possible that salicylate acts by decreasing ATP levels near K_ATP_ channels or activating AMPK. Moreover, high doses of salicylate potentiate the depolarization-induced suppression of excitation (DSE) on the excitatory synapses on the CW neurons (Zugaib and Leão, 2018), and, interestingly, activation of AMPK has been shown to potentiate DSE on hypothalamic neurons (Borgquist et al., 2015). The effects of salicylate on the cartwheel neuron can have a profound impact on the excitability of the fusiform neurons, and K_ATP_ channel regulation may be one important factor contributing to the development of tinnitus by this drug.

## Acknowledgments

Supported by FAPESP grants (2016/01607-4 and 2019/13458-1). We thank Mr. J. Fernando Aguiar for the technical support, Dr. Henrique von Gersdorff for insightful discussions about the data and Dr. Christopher Kushmerick for reviewing the manuscript.

